# Fast and efficient *Borrelia* genome recovery from tick samples using Whole-Genome Amplification

**DOI:** 10.64898/2025.12.02.691817

**Authors:** Sophie Melis, Cristina Cezara Bunduc, Laura Mottarlini, Michela Vumbaca, Irene Mileto, Sabrina Hepner, Gabriele Margos, Volker Fingerle, Andreas Sing, Paola Prati, Claudio Bandi, Vittorio Sambri, Stefano Gaiarsa, Fausto Baldanti, Greta Bellinzona, Davide Sassera

## Abstract

*Borrelia burgdorferi* sensu lato bacteria are the causative agents of Lyme borreliosis, a multisystemic illness with an expanding epidemiology in temperate areas. Genomic studies on *Borrelia* are hindered by the difficulty of culturing procedures: standard protocols require extended incubation, substantial technical expertise, and are prone to failure, limiting timely recovery of isolates. This contributes to a low number of complete *Borrelia* genomes available in public repositories, particularly for species other than *B. burgdorferi* sensu stricto.

Here we introduce a novel approach that overcomes the necessity for extended culture by utilizing Whole Genome Amplification (WGA) directly on freshly collected ticks, and that can be performed in parallel to classical culturing. The protocol is paired with a tailored bioinformatic pipeline designed to ensure accurate assembly and reliable downstream analyses. Benchmarking on multiple control isolates demonstrated that the method yields high-quality chromosomal assemblies. To demonstrate practical applicability, we applied the protocol to freshly collected ticks, successfully generating five high-quality *Borrelia* chromosomes (two *B. lusitaniae*, two *B. afzelii* and one *B. garinii*). By generating sequencing-ready DNA in five days rather than months, our protocol greatly streamlines the process and minimizes the effort associated with traditional culture-based methods. This workflow will facilitate broader representation of understudied *Borrelia* species and support future epidemiological, ecological, and evolutionary investigations on this pathogen.

## Introduction

Bacteria of the *Borrelia burgdorferi* sensu lato (s.l.) complex are the causative agents of Lyme borreliosis (LB), the most prevalent vector-borne illness in the Northern hemisphere, with around 30,000 confirmed cases in the United States and around 85,000 reported cases in Europe each year ^1–3^. A typical early sign of LB is an expanding skin lesion known as erythema migrans that, if left untreated, can progress to a disseminated infection. This may develop into severely debilitating stages characterized by arthritis, neurological disorders, carditis and acrodermatitis chronica atrophicans ^4^.

The *B. burgdorferi* s.l. complex currently comprises 28 validated and *Candidatus* species^5^. These different species, and even different isolates within the same species, can cause different clinical manifestations ^3,6^. For instance, *B. garinii* and *B. bavariensis* are more associated with neuroborreliosis, while *B. afzelii* mainly causes skin infections ^4,7^. The genetic bases of such differences are currently unclear. In Europe, the primary vector is the hard tick *Ixodes ricinus* and the most common circulating *Borrelia* species include *B. afzelii, B. garinii, B. burgdorferi* sensu stricto (s.s.), *B. valaisiana*, and *B. lusitaniae* ^8^. While the first three species are known human pathogens, no clinical cases have been reported for *B. valaisiana* (Margos et al. 2017), and the pathogenic potential of *B. lusitaniae* remains poorly understood ^9,10^.

Bacterial genomics now is a key tool in microbiological research and allows to refine phylogenetic reconstructions, to improve epidemiological characterizations, and to discover genomic determinants of pathogenicity, virulence, and resistance to antimicrobial agents. Unfortunately, despite several notable efforts ^11–14^, *Borrelia* genomics have been lagging behind compared to other pathogens, with relatively few high-quality genomes publicly available for species outside *B. burgdorferi* s.s. ^15^.

The unusual genome architecture of *Borrelia*, which combines a linear chromosome with up to 21 linear and circular plasmids (ranging from 5–84 kbp in size) represents a challenge for genomic studies ^11,16^. Additionally, an even greater challenge lies in the difficulty of obtaining pure cultures. Isolation from patients typically relies on tissue biopsies or aspirates, an invasive procedure that is rarely recommended and not always feasible. Culturing from ticks or patient material is inefficient: it requires one to two months of incubation depending on the species, and in our experience yields successful isolates from only about 25% of PCR-positive samples. Moreover, culture-based approaches are not only slow but also labor-intensive and resource-demanding, limiting their scalability for large epidemiological or comparative genomics studies. As a result, many *Borrelia* isolates risk never being sequenced, leaving important gaps in our understanding of the diversity and pathogenic potential of this genus. These drawbacks highlight the need for alternative strategies to obtain *Borrelia* genomes without relying solely on culture.

Here, we describe a novel workflow that combines Whole Genome Amplification (WGA) with dedicated bioinformatic analyses, enabling the recovery of sequencing ready *Borrelia* samples from single ticks within five days. WGA allows amplification of genomic DNA from very small starting quantities, bypassing the need for long culture periods and enabling genome sequencing from early-stage cultures or low-input samples. In fact, as little as picograms of starting DNA is sufficient for WGA, compared to the nanograms required for genome sequencing even with the more sensitive Illumina library preparation protocols. Our approach shows substantially higher success rates compared to conventional methods, while remaining compatible with parallel PCR-based diagnostics and classical culturing procedures.

## Materials and Methods

### Overview of the experimental design

Two approaches were used for the generation of *Borrelia* genomic DNA: a standard culturing protocol (hereafter: standard protocol) ^11^ and a novel protocol developed by us that leverages whole genome amplification (hereafter: WGA protocol) (Figure 1). Three pure cultures of *Borrelia* isolates, previously obtained from ticks and stored in our collection, were processed with both protocols. These samples are hereafter referred to as “positive pure culture controls” and were used to compare the quality of the genomes generated with both protocols.

**Figure 1.**
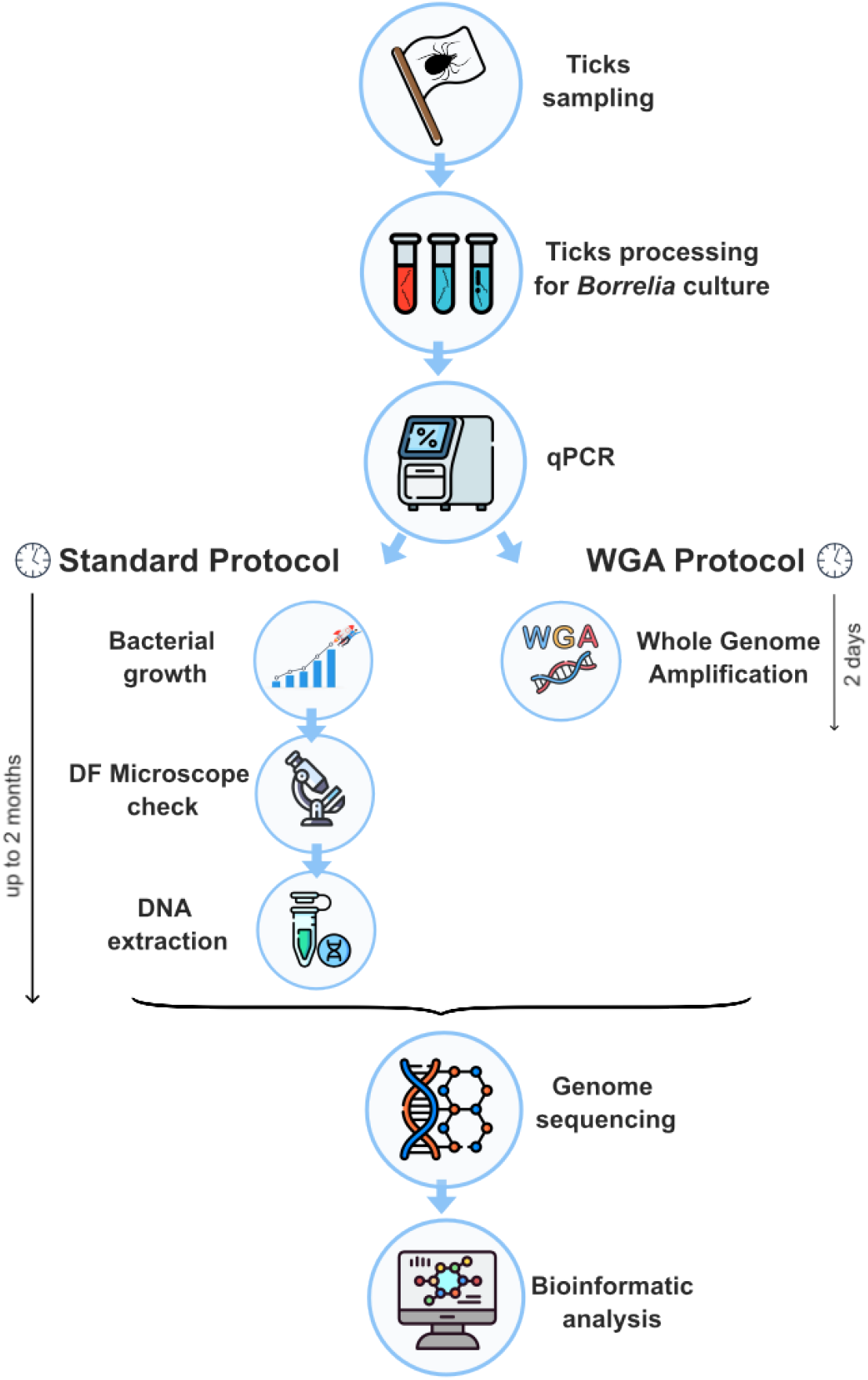
Comparison between standard protocol and the newly optimized WGA protocol.

### Samples collection and culturing conditions

We used three pure culture controls of *Borrelia* spp. previously isolated from *I. ricinus* ticks between 1989 and 1996 in northern Italy which had undergone approximately 13 culture passages. The cultures, stored at -80°C in the laboratory of the National Reference Center for *Borrelia* (Oberschleissheim, Germany), were revived using theModified Kelly-Pettenkofer (MKP) medium, supplemented with 6% of rabbit serum at 33°C, in 8 ml glass tubes ^17^.

In parallel, a total of 26 ticks (nymphs and adults) were collected via dragging in multiple locations in northern Italy and morphologically identified as *I. ricinus* (see Supplementary Table S1 for collection details). Single individuals were then processed by surface sterilization, cut with a sterile scalpel and cultured in 1.5 ml plastic tubes with 350 μl of MKP medium, supplemented with antibiotics (Sulfamethoxazole 45 μg/ml, Fosfomicin 170 μg/ml and Amicacin 25 μg/ml) ^17^.

### Molecular processing - qPCR and Whole Genome Amplification (WGA)

Positive culture controls were cultured for 7-14 days and examined daily under dark field microscope (Nikon - ECLIPSE) to check the concentration and viability of spirochetes, until they reach a concentration of 10^7^ per ml. Volumes of 20 ml of positive cultures were centrifuged at 4500 rpm for 20 minutes. The pellet was recovered and DNA extraction was performed using the NucleoSpin Tissue Kit (Macherey-Nagel) according to the manufacturer’s instructions. In parallel, a 150 μl volume of each culture was centrifuged at 8000 rpm for 10 minutes and the pellet resuspended in 4 μl of PBS. WGA-protocol was then performed according to the manufacturer instructions (REPLI-g Single Cell Kit, Qiagen).

Newly collected ticks were subjected to culture protocol as described above. Three days after tick processing, a volume of 2 μl of each new culture was collected and subjected directly to qPCR, to assess the presence of *Borrelia*. Amplification of a fragment of 23S rRNA gene, was performed using real-time PCR with the following thermal protocol: 95°C for 5 min, 40 cycles at 95°C 15s and 60°C 30s, and melt curve from 55°C to 95°C with increasing increments of 0.5°C per cycle (Michelet et al. 2014). PCR-positive cultures were then checked under dark-field microscopy. Cultures were determined as PCR-positive if specific amplification was observed with a threshold cycle <32.

A second aliquot of 150 μl of medium of qPCR-positive samples was collected and subjected to the optimized WGA-procotol. First, the aliquot was centrifuged at 8000 rpm for 10 minutes. Then, to evaluate the separation of *Borrelia* from tick DNA in the 150 μl aliquot from the culture, after centrifugation, both pellet and supernatant were subjected to qPCR specific for *Borrelia* (23S rRNA gene) and a tick housekeeping gene (*calreticulin*) ^18,19^. WGA was then performed as above. A control qPCR for *Borrelia* (same condition as before) was performed after WGA protocol, and the samples with a threshold cycle <20 were further processed for sequencing.

### Whole Genome Sequencing (WGS)

All DNA samples (DNA from positive controls from the standard protocol, DNA from WGA of positive controls and fresh tick samples) were subjected to DNA quantification using Qubit™ (dsDNA Quantification Assay, Broad-range) and followed by Nanopore and Illumina sequencing. Standard Library Preparation was performed for Illumina sequencing using Illumina DNA Prep following manufacturer instructions (final concentration of each sample was 0.2 ng/μl). Libraries were then sequenced on a MiSeq machine and v2 (500 cycles) reagents multiplexing in order to obtain 1Gb per sample. In parallel, aliquots of the same DNA samples were subjected to Nanopore sequencing (outsourced to the company Eurofins, using Oxford Nanopore GridION, flow cell R10.4.1, using v14 library prep chemistry, declaring a minimum of 210 Mb of sequencing data per sample).

### Read Quality Control

Illumina and Nanopore reads were quality-checked using FastQC v0.12.1 and summarized using MultiQC v1.23 ^20^. Adapters and low-quality bases (Q < 20) were trimmed in Illumina reads using fastp v1.0.1 ^21^.

All quality-filtered reads were then classified using Kraken2 v.2.1.16 ^22^ with the `core_nt` database. This step was used both to identify and remove potential contaminant reads (i.e., non-*Borrelia* taxa), and to confirm the taxonomic identity of the *Borrelia* species present in each sample. KrakenTools ^23^ `*extract_kraken_reads*.*py*` script was employed to retrieve reads assigned to the *Borrelia* genus (taxid: 64895) and its subordinate taxa, ensuring that only genus-specific sequences were retained for downstream analyses.

### Genome Assembly and Quality Assessment

Hybrid genome assemblies were generated using Unicycler v.0.5.1 ^24^, which integrates short Illumina reads and long Nanopore reads for improved assembly. Unicycler was run with default parameters in bold mode. Assemblies were polished within Unicycler using both read types and manually curated. Assembly quality was then assessed using BUSCO v.6.0 ^25^, using the Spirochaetia lineage dataset, to evaluate completeness. Contamination was assessed using CheckM2 v.1.0.2 ^26^. Assemblies generated from the same sample using different protocols (standard vs WGA) were compared using blastn within Bandage v.0.8.1 ^27^.

### Plasmids identification and typing

Plasmids were first identified in the assembled genomes through manual inspection using Bandage v.0.8.1 ^27^. Plasmid nomenclature was assigned according to the paralogous gene families (PFam) they contained, specifically PFam32, PFam49, PFam50, and PFam57/62 ^16,28–31^. The identification process followed the general approach outlined by Hepner et al. (2023), using the same reference PFam set of sequences. To detect PFam32 plasmid partitioning gene loci within the assembled contigs, we employed BLASTN v.2.16.0 ^32^ using default parameters. Loci showing at least 86.5% nucleotide identity and a minimum of 90% alignment coverage were considered valid matches and used to assign plasmid designations accordingly.

### Typing

Multi-locus sequence typing (MLST) and core genome MLST (cgMLST) typing were performed using the assembled genomes of each isolate as input, following the published schemes ^33–35^. Assemblies were uploaded to the PubMLST database for *Borrelia* spp. (https://pubmlst.org/) ^36^, where MLST and cgMLST alleles, sequence types (STs) and core genome sequence types (cgSTs) were automatically assigned. A minimum spanning tree (MST) was generated using GrapeTree v.1.5.0 ^37^, as implemented in the PubMLST platform ^36^ using the MLST profiles. MLST profiles were retrieved for all available *Borrelia* genomes (n = 519, 8-locus scheme) and combined with the eight novel isolates sequenced in this study. The MST was constructed using the “MSTree V2” algorithm with default parameters.

### Read Mapping and Coverage Analysis

To assess whether the amplification and sequencing process introduced any systematic coverage biases across the chromosome, raw Illumina reads were mapped for each sample to the reference genome of the *Borrelia* species identified by Kraken2 in each sample, using Bowtie2 v.2.5.4. The reference genomes used were: *B. garinii* (CP075232.1) and *B. burgdorferi* s.s. (CP161191.1). Mapping quality and coverage metrics were assessed using Samtools v1.21 ^38^. Coverage uniformity plots were created using a custom Python script.

## Results and Discussion

We designed a lab protocol based on Whole Genome Amplification (WGA) to be performed in parallel to the standard protocol for *Borrelia* isolation in culture to obtain genomes faster, and with higher success rate. To validate the approach we first evaluated the quality of WGA genomes from pure cultures. Then, we applied this novel protocol and the associated bioinformatic pipeline to fresh tick samples.

### WGA allows to obtain accurate Borrelia genome assemblies

To assess the performance and reliability of the whole-genome amplification (WGA) approach, we first evaluated its impact on genome quality, testing it on three *Borrelia* positive pure culture controls, specifically one *B. burgdorferi* s.s. and two *B. garinii* samples. DNA was extracted from these cultures and subjected both to WGA and to standard DNA extraction workflow and then sequenced using Illumina and Nanopore platforms.

We initially assessed the uniformity of coverage by comparing the two approaches across the three samples. As expected, coverage plots from the WGA protocol exhibited a more uneven distribution across the chromosome compared to the smooth, uniform coverage obtained with the standard protocol (Figure S1). This pattern reflects the well-documented amplification biases inherent to WGA methods. Importantly, however, no chromosomal regions were left uncovered in WGA assemblies. This indicates that, despite minor biases, the WGA approach provides comprehensive chromosomal representation suitable for downstream analyses.

Using the same hybrid assembly method for both standard and WGA-derived datasets, we generated complete genome assemblies for the three *Borrelia* samples and subsequently compared their quality and completeness (Table 1). In both *B. garinii* isolates, IT-G1 and IT-G2, the chromosome was assembled into a single contig both from pure culture and from WGA. For IT-B1, a complete chromosome assembly was obtained only with the WGA protocol, while the standard approach produced a more fragmented result. Overall, chromosomal sequences were nearly identical across methods, with only minimal SNP differences (≤4 per genome) and comparable completeness scores (BUSCO: 98–100%; CheckM2: 92.85–96.44%). Assemblies from WGA samples tended to be slightly smaller, with repeated telomeric regions not fully reconstructed—as observed in IT-B1—and fewer plasmids recovered (3– 5 vs. 6–9), occasionally showing structural rearrangements such as plasmid– chromosome fusion (e.g., lp36 in IT-G1).

**Table 1.** Comparison of WGA and standard protocol for three pure culture samples. ND stands for not determined. * Independent linear or circular sequences. £ This plasmid was fused to the chromosomal telomere by the assembly and after inspection was manually separated.

|  | Standard Protocol | WGA Protocol | Standard Protocol | WGA Protocol | Standard Protocol | WGA Protocol |
| --- | --- | --- | --- | --- | --- | --- |
|  | IT-B1 |  | IT-G1 |  | IT-G2 |  |
| Species | *B. burgdorferi* s.s. |  | *B. garinii* |  | *B. garinii* |  |
| Chromosomal contigs | 8 | 1 | 1 | 1 | 1 | 1 |
| N. of plasmids* | 9 | 3 | 6 | 5 | 7 | 4 |
| Plasmids | lp28-2,<br>lp36,<br>cp26, | cp32-3<br>(partial),<br>cp32-7 | lp54,<br>lp28-7,<br>lp36, | lp54,<br>lp36<sup>£</sup>,<br>cp26, | lp54,<br>lp36,<br>lp17, | cp32-1,<br>cp32-5,<br>cp32-6, |

|  | cp32-1,<br>cp32-3+7,<br>cp32-5,<br>cp32-9,<br>cp32-11,<br>cp32-12 | (partial),<br>cp32-11<br>(partial) | cp26,<br>cp32-4,<br>cp32-9 | cp32-4,<br>cp32-9 | cp32-1,<br>cp32-5,<br>cp32-6,<br>cp26 | cp26 |
| --- | --- | --- | --- | --- | --- | --- |
| Chromosome Size (bp) | 951,498 | 908,642 | 905,758 | 905,754 | 906,465 | 906,465 |
| N50 | 213,238 | 908,642 | 905,758 | 905,754 | 906,465 | 906,465 |
| Completeness % (BUSCO) | 100 | 98 | 98 | 98 | 99,7 | 99,7 |
| Completeness % (CheckM2) | 96,44 | 93,9 | 93,26 | 92,85 | 94,99 | 94,11 |
| Contamination % (CheckM2) | 0,27 | 0,27 | 0,48 | 0,48 | 0,44 | 0,43 |
| ST | 20 | 20 | 187 | 187 | 177 | 177 |
| cgST | 440 | 440 | 444 | 444 | 438 | 438 |

Plasmid assembly proved particularly challenging across all samples and methods. In the three culture controls, plasmids such as lp54, cp26, lp17, lp36, lp28-2, lp28-7, and multiple cp32 variants were detected. WGA assemblies recovered only a subset of plasmids, with some appearing incomplete or absent. Notably, lp54 is considered constitutive ^28,29^, but was missing in IT-B1 for both protocols. The fragmentation of the IT-B1 chromosome observed with the standard protocol, along with the absence or partial recovery of several plasmids, likely results from a combination of factors. These could include biological and culture-related issues such as long-term storage, multiple passages (13), and potential DNA degradation together with technical limitations in the assembly process, including difficulties in resolving repetitive or low-coverage regions with the assembler, and the naturally low copy number of *Borrelia* plasmids ^39^.

### Borrelia genomes from fresh tick samples

We then applied our protocol to newly collected ticks. 26 specimens of *I. ricinus* were collected, sterilized and put in MKP medium for *Borrelia* isolation. After 3 days, all samples were tested for *Borrelia* using qPCR, and positives (7 samples out of 26, 27%) were centrifuged, a process that efficiently enriched *Borrelia* cells in the pellet allowing us to retrieve a sufficient amount of DNA for genome amplification (Supplementary Table S2 - sheet 1 and 2). These samples were thus subjected to WGA and sequencing with Illumina and Nanopore technologies.

While long-term pure cultures provided high-quality *Borrelia* DNA with minimal contamination, samples obtained from early cultures derived directly from tick material and processed using WGA inevitably contained co-amplified non-*Borrelia* DNA. Such contaminations can complicate downstream genomic analyses, leading to biased assemblies or erroneous conclusions if not properly accounted for. To address this issue, we developed an ad-hoc bioinformatic pipeline. We first used Kraken2^22^ to perform taxonomic classification. In the three long-term pure cultures, Kraken2 ^22^ classified over 97% of reads as *Borrelia*, confirming the high purity of these samples. However, fresh tick samples amplified using WGA exhibited variable levels of contamination of reads assigned to non-*Borrelia* taxa (Supplementary Table S4). Within these, the proportion of tick-derived reads was consistently low, never exceeding 2.5% of total reads assigned to the Ixodidae family, confirming that centrifugation allows efficient separation of bacterial cells from tick DNA. Most of the contaminants are represented by other bacteria, known symbiont and tick-borne pathogens, including *Spiroplasma*^40^, *Midichloria*^*41*^ and *Rickettsia*^42^. Furthermore, Kraken2 classification provided species-level confirmation of *Borrelia* isolates across all samples, validating the identity of both standard and WGA-derived datasets.

Two out of seven samples, which showed a high contamination level (reads assigned to the *Borrelia* genus <30 Mb; <1% of total), were excluded from downstream analyses. Remarkably, sample IT-L3, showing only 4.73% of reads assigned to *Borrelia*, still achieved 130X coverage, and was thus maintained. The full Kraken classification results for all samples can be found in Supplementary Table S4.

All non-*Borrelia* reads identified by Kraken2 were removed prior to downstream bioinformatics steps. After filtering, the retained *Borrelia* reads were successfully assembled to nearly complete and contiguous genomes with contigs number ranging from 4 to 52 and N50 values spanning from 129,633 to 905,836 bp, reflecting good contiguity (Table 2). Notably, four out of five genomes (IT-A1, IT-A3, IT-L3, IT-G3) reached chromosome-level resolution, with a completely reconstructed linear chromosome and several plasmid sequences recovered.

**Table 2.**
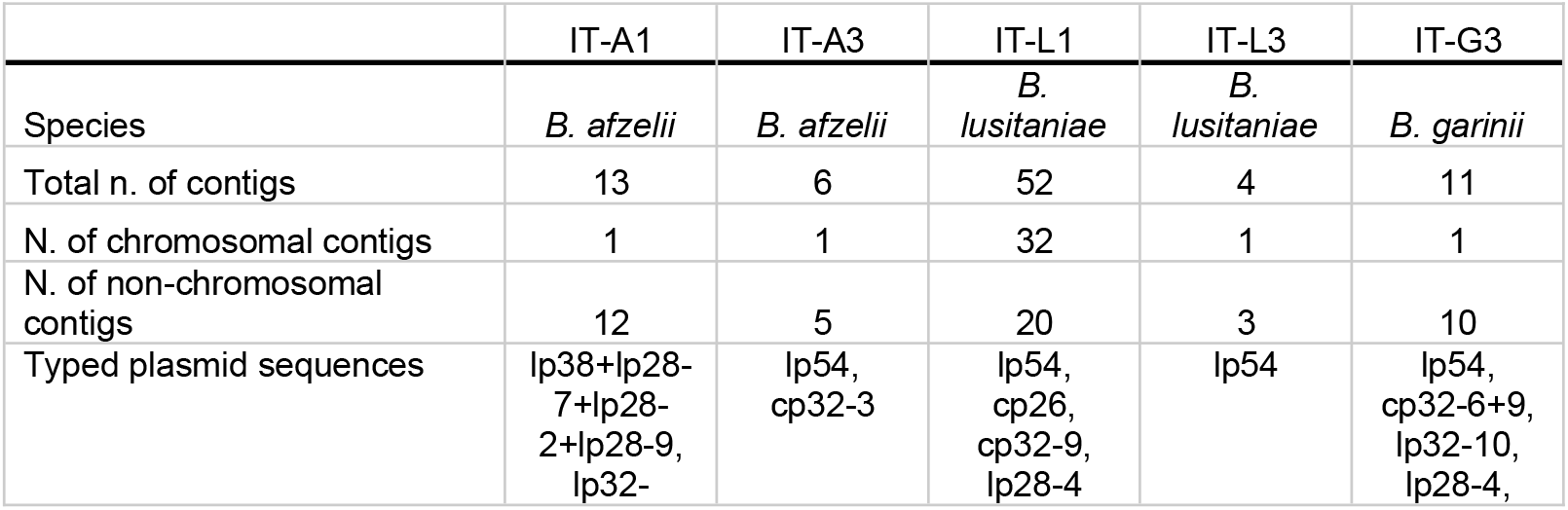
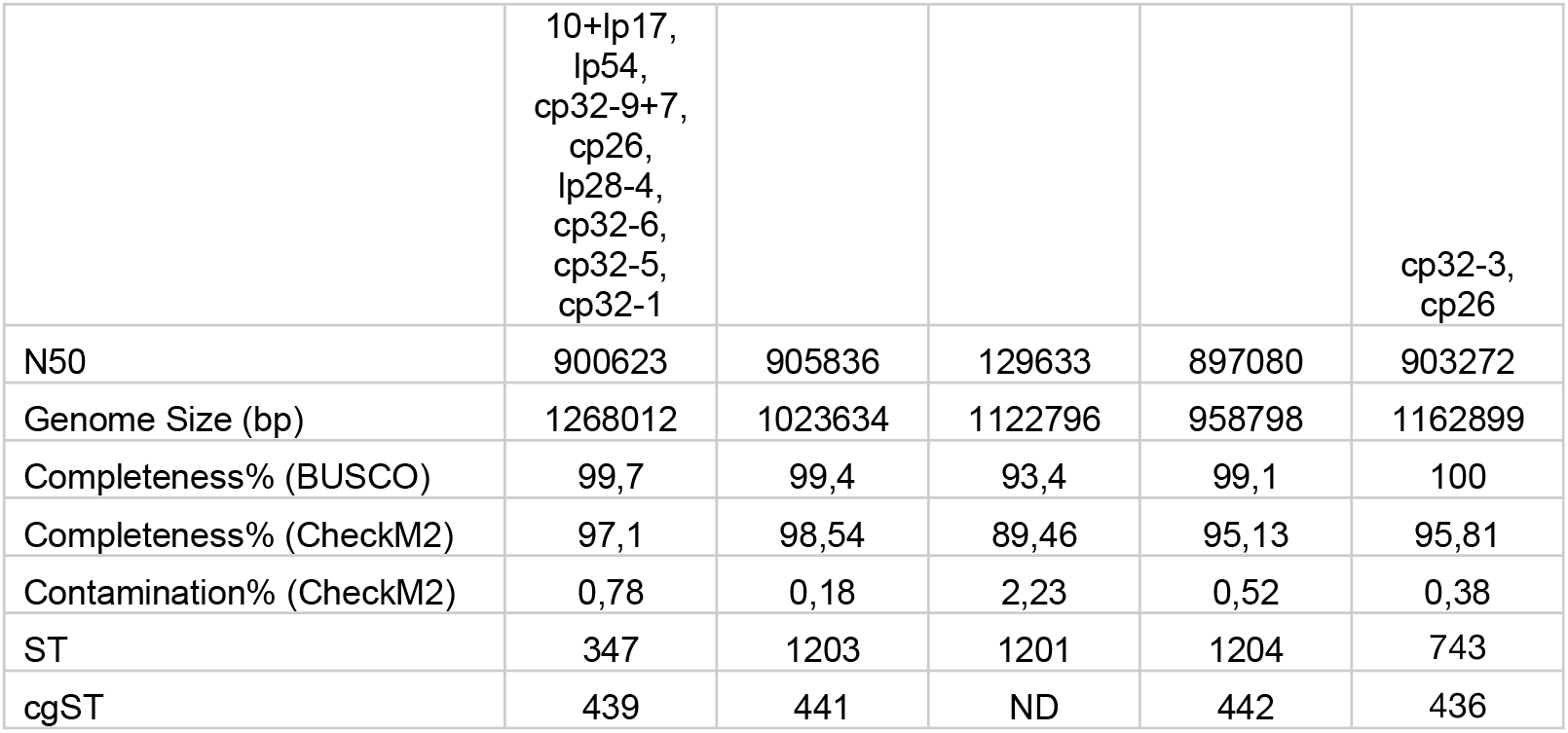
Assembly statistics, quality metrics, and typing results for the five samples processed exclusively with the WGA protocol.

Plasmids were typed based on the presence of PFam32, PFam49, PFam50, and PFam57/62 following established literature. Assignment to a known plasmid type was performed only when a PFam match was detected. Many contigs carrying partial plasmid sequences remained untyped. Some of these untyped contigs could represent previously uncharacterized plasmids, warranting future dedicated investigation, while others may simply be incomplete due to missing PFam domains, reflecting the known limitations of our assembly and typing approach for *Borrelia* plasmids. Completeness of plasmid assemblies varied due to several factors, including sequence similarity among cp32 variants, structural complexity, and differences in copy number, which can hinder full reconstruction even from high-quality assemblies. Plasmid recovery varied between isolates, but lp54 was recovered as a complete plasmid in all five genomes, consistently with reports on its stability^28,29^. Similarly, cp26, another constitutive plasmid, was detected in three out of five isolates, possibly due to limitations in plasmid recovery using WGA protocol.

### WGA genomes allow in depth downstream analyses

To assess the genotypic diversity of the sequenced isolates, we first performed MLST using the standard 8-locus *Borrelia* scheme ^43,44^. All samples, comprising both positive controls and fresh samples were successfully typed, resulting in the assignment of three novel sequence types (STs) for three fresh samples IT-A3, IT-L1, and IT-L3 (Table 2).

To further contextualize our isolates within the global diversity of *B. burgdorferi* s.l. complex, we built a minimum spanning tree (MST) based on all the complete MLST profiles associated with a sample deposited on *Borrelia* PubMLST. The MST (Figure 2) shows the overall population structure of *B. burgdorferi* s.l., with clear species-level clustering. The two *B. afzelii* isolates (IT-A1, IT-A3) were embedded within the main *B. afzelii* cluster (colored in light orange in Figure 2), in line with the species assignment. Isolate IT-B1 also grouped within the central cluster corresponding to its species, *B. burgdorferi* s.s. (colored dark blue in Figure 2). The *B. lusitaniae* isolates (IT-L1, IT-L3, colored dark green in Figure 2) formed a divergent point and a peripheral branch, reflecting their distinct genetic background. The *B. garinii* isolates, IT-G1, IT-G2, and IT-G3, clustered within the species *B. garinii*.

**Figure 2.**
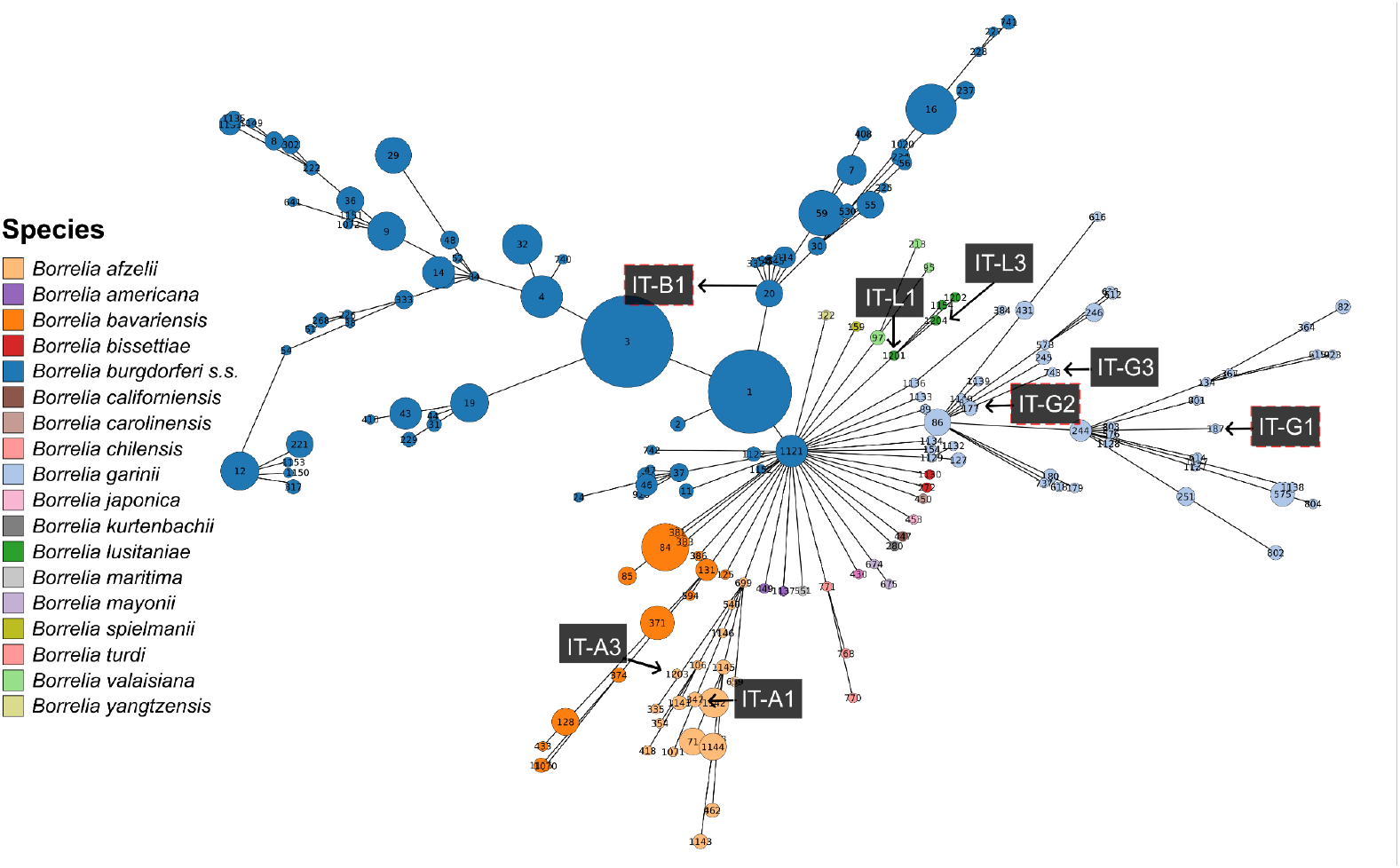
Minimum spanning tree of *Borrelia burgdorferi* s.l. complex based on 8-locus MLST profiles. The tree was constructed with the built-in BIGSdb GrapeTree tool using all available sequence types associated with a genome and a complete MLST profile (n = 521), together with the eight isolates sequenced in this study. Each node represents a sequence type (ST), with its size proportional to the number of isolates corresponding to the ST. Colors correspond to species as indicated in the legend. Edges connect STs according to allelic differences. Isolates from this study are highlighted with black boxes and labels, with an additional red dashed border for positive controls (IT-B1, IT-G1, IT-G2).

We subsequently performed cgMLST analyses ^45^. In contrast to MLST, which relies on a few housekeeping loci (eight MLST loci for *Borrelia*), cgMLST exploits several hundred conserved loci across the genome, enabling much finer resolution (the *Borrelia* cgMLST scheme contains 639-loci). All samples were assigned to novel cgMLST types, with the exception of one *B. lusitaniae* isolates (IT-L1) where cgST assignment failed. Initially, for both *B. lusitaniae* isolates IT-L1 and IT-L3, cgMLST assignment failed. In this case, the loci were present but were too divergent to meet the established thresholds (99% query coverage and 97% identity) for allele assignment in cgMLST. This outcome reflects the fact that *B. lusitaniae* was not included in the development genome set of the cgMLST scheme ^45^ as high quality genomes were not available for this species by this time. After seeding the *Borrelia* PubMLST database by some manual curation including assigning exemplary alleles that were slightly different in lengths to those of the previously included alleles, the cgMLST scheme performs well for the species *B. lusitaniae* and one out of the two samples, IT-L3, was successfully assigned to a cgST. IT-L1 was not assigned to a cg-ST due to the presence of only 590 loci out of 639, likely reflecting an incomplete genome, as also suggested by its relatively low BUSCO score (93.4%).

The previously absence of high-quality *B. lusitaniae* genomes in the *Borrelia* PubMLST and other databases underlines the understudied status of this species and the need for broader genomic representation of neglected species.

## Conclusions

In this study, we developed and validated an optimized workflow integrating whole genome amplification (WGA) in the standard *Borrelia* culturing procedures, to enable faster, easier, and more successful genome sequencing. Overall, our WGA-based protocol allows a substantial improvement in the success rate and reduces the sample-to-sequencing timeline. Whereas standard protocols require 30–60 days of culture and frequently fail to yield pure isolates, the WGA-based workflow can be completed in just 5 days, producing sequencing-ready samples.

Our results demonstrate that WGA enables the recovery of high-quality chromosome assemblies even from low-input samples or early-stage cultures, overcoming a major bottleneck in generating genomic data for this difficult to grow bacterium. While WGA-derived assemblies yield highly accurate and complete chromosomal reconstructions, plasmid recovery is often incomplete due to intrinsic sequence complexity, high similarity among cp32 variants, and amplification biases associated with WGA. Despite this limitation, the chromosomal information generated is sufficient to support a broad range of genomic epidemiology applications, including sequence typing (MLST and cgMLST), comparative genomics, and studies of strain diversity, population structure, and geographic distribution. Crucially, this strategy makes it possible to recover genomic data from isolates with limited success of culturing (25% of qPCR positive cultures), and it has already enabled the identification of novel sequence types, expanding the representation of under-sampled species such as *B. lusitaniae* in public databases.

## Data Availability

All sequencing reads and genome assemblies generated in this study have been deposited in NCBI under BioProject PRJNA1330164 and submitted to PubMLST. Sample-specific accession numbers, PubMLST identifiers, and relevant metadata are provided in Supplementary Table S3. All scripts and curated plasmid reference datasets generated in this study are publicly available on GitHub at https://github.com/MIDIfactory/Borrelia-WGA-Project.

## Acknowledgment

This work was supported by the grant Ricerca corrente 2024 from Fondazione IRCCS Policlinico San Matteo to Davide Sassera. We would like to acknowledge Keith A. Jolley, the developer of BIGSdb, for his support in manually curating and seeding the database, which enabled the *Borrelia* cgMLST scheme hosted on the *Borrelia* PubMLST website to be applied for the species *B. lusitaniae*.

