## Supplementary Figure S1 for "Fast and efficient *Borrelia* genome recovery from tick samples using Whole-Genome Amplification"

### Supplementary Figures

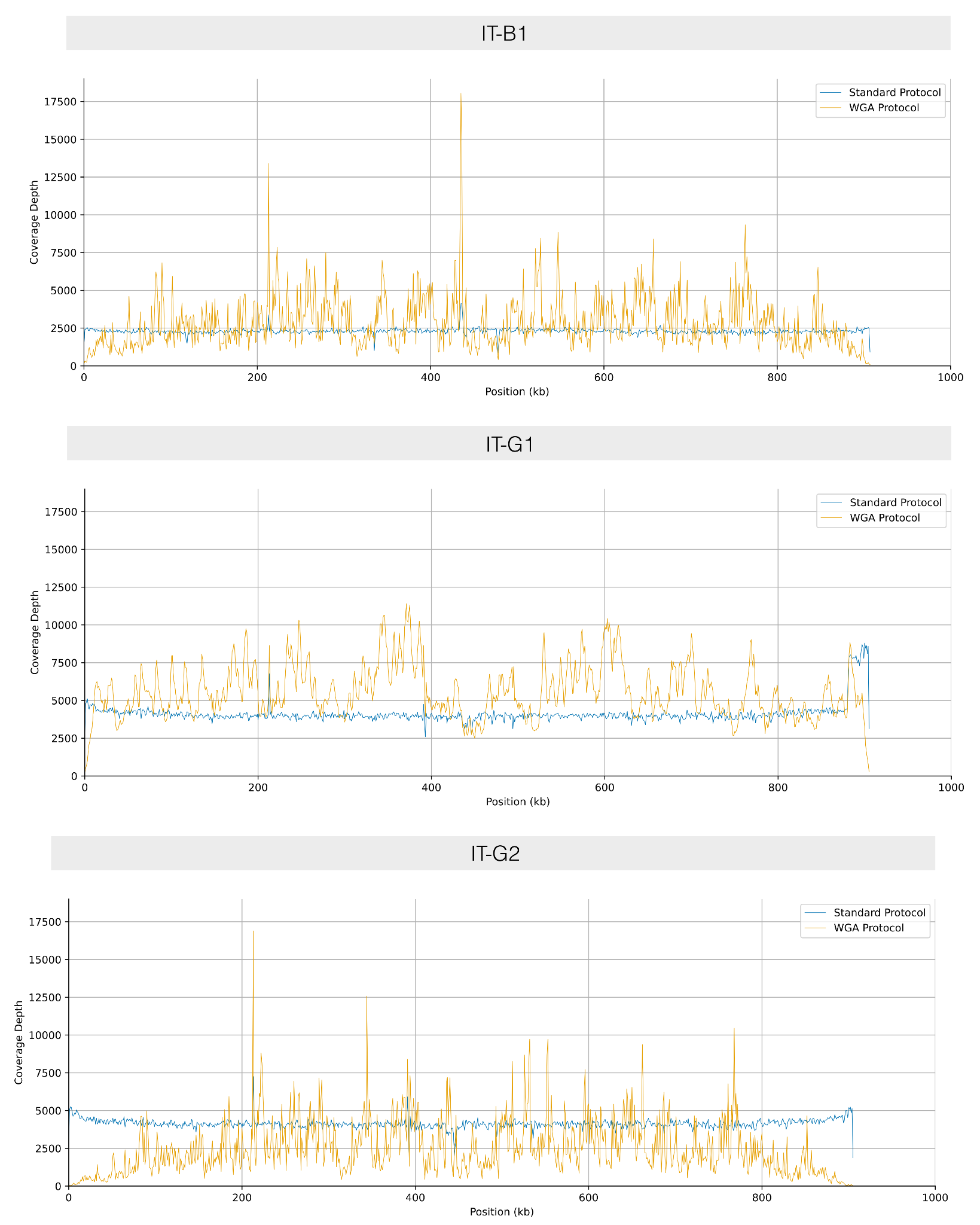

**Supplementary Figure S1**. Coverage profiles of the three pure culture controls (IT-B1, IT-G1, IT-G2) processed with standard and WGA protocols. Average sequencing depth was calculated in non-overlapping 1 kb windows along the reference genome. For each isolate, coverage obtained using the standard sequencing protocol (blue) is compared with coverage obtained using whole-genome amplification (WGA, orange).
